# Single-cell foundation models reveal context-sensitive cancer programmes under subtype shift

**DOI:** 10.64898/2026.04.28.721114

**Authors:** James Wallace, Gehad Youssef, Namshik Han

## Abstract

Single-cell foundation models (scFMs) have shown promise as transferable representations of cellular state, but recent zero-shot evaluations suggest that they do not consistently outperform simpler baselines. We asked whether this apparent limitation reflects an intrinsic weakness of scFMs or instead the difficulty of using them without task-specific adaptation. To test this, we fine-tuned two widely used scFMs, Geneformer and scGPT, on common tumour subtypes from renal, lung, and breast cancer, and compared them with a LightGBM baseline on within-domain validation cohorts and on out-of-domain rarer, unseen cancer subtypes. Across all three organs, the models achieved near-perfect within-domain discrimination (AUROC 0.98-1.00), but differences emerged under subtype shift. On chromophobe RCC, scGPT and Geneformer achieved AUROC 0.88 and 0.92 respectively versus 0.64 for LightGBM; on SCLC, Geneformer reached 1.00 versus 0.82 for LightGBM; and on TNBC, scGPT achieved 0.80 versus 0.49 for

LightGBM. To determine whether this generalisation reflected meaningful adaptation rather than arbitrary feature drift, we applied Integrated Gradients, an interpretability technique, to the fine-tuned scFMs and SHAP to LightGBM. LightGBM showed highly stable gene-importance rankings across datasets, whereas the foundation models were substantially more context-sensitive. However, this flexibility was not random: all models converged on a shared within-domain core, while scFMs acquired larger rare-subtype-specific gene sets and pathway programmes during transfer. Pathway enrichment further supported the biological relevance of these attributed genes. Together, these results suggest that fine-tuned scFMs can bridge clinically relevant domain shifts in cancer single-cell analysis and that interpretability provides a practical route to distinguishing biologically grounded adaptation from rigid reuse of training-era rules.

## Introduction

Foundation models trained on millions of single cells have reframed transcriptomics as a transfer learning problem. Models such as scGPT [1] and Geneformer [2] adapt the transformer architecture [3] to learn generalisable representations of cellular states from large-scale transcriptomic data [4, 5]. Despite their theoretical promise, scFMs have recently faced significant scrutiny. Several independent benchmarks have demonstrated that in zero-shot settings, these computationally expensive models frequently fail to outperform simple linear baselines or traditional machine learning approaches across a variety of downstream tasks [6, 7, 8]. This raises a critical question: is the high computational cost of pre-training and deploying scFMs justified by their performance?

While zero-shot evaluation highlights current limitations, recent evidence suggests that parameter-efficient fine-tuning can unlock the true potential of scFMs [9, 10]. Indeed, we have also found that simpler, less costly baselines such as LightGBM [11] are exceedingly difficult to outperform at cell type classification when applied within-domain to the same task that was trained on. However, the true test of a foundation model lies not in its ability to interpolate within a familiar distribution, but in its capacity for domain adaptation: the ability to align and transfer knowledge across heterogeneous datasets [12].

Domain adaptation is particularly crucial in oncology, where clinical decision-making often concerns rare subtypes, metastatic lesions, or small patient cohorts for which labelled data are scarce [13]. Rare cancer subtypes often lack sufficient data to train robust, subtype-specific machine learning models from scratch [14]. Transfer learning approaches have shown promise in overcoming these data limitations by exploiting molecular patterns shared across cancers [15, 16]. If foundation models are useful here, they should transfer compact cancer-relevant programmes from common tumour types to rarer subtypes more effectively than conventional machine learning baselines.

We therefore investigated domain adaptation in a controlled cell-state classification setting spanning three organs. Models were trained on common tumour subtypes: clear cell renal cell carcinoma (ccRCC), lung adenocarcinoma (LUAD), and oestrogen receptor-positive (ER+) breast cancer, and then evaluated both on held-out cohorts of the same subtype and on rarer out-of-domain contexts: bone metastatic and chromophobe renal cancer, lung squamous cell carcinoma (LUSC) and small cell lung cancer (SCLC), and HER2+ and triple-negative breast cancer (TNBC). LightGBM was used as a strong classical baseline because it is inexpensive, interpretable, and often difficult to beat for supervised tabular classification [11].

Beyond performance, we asked how each model adapts. Because attention weights are frequently used to interpret transformer models but are often not faithful explanations of model behaviour [17], we used feature-attribution methods that act directly on the prediction function: Integrated Gradients (IG) [18] for the fine-tuned scFMs, a robust gradient-based attribution method [19], and SHAP for LightGBM [20]. This made it possible to compare not only whether models generalised, but also whether they did so by reusing rigid gene rules or by reweighting biologically plausible programmes in a context-dependent manner. A schematic overview of the complete analytical pipeline is presented in Figure 1.

**Figure 1.**
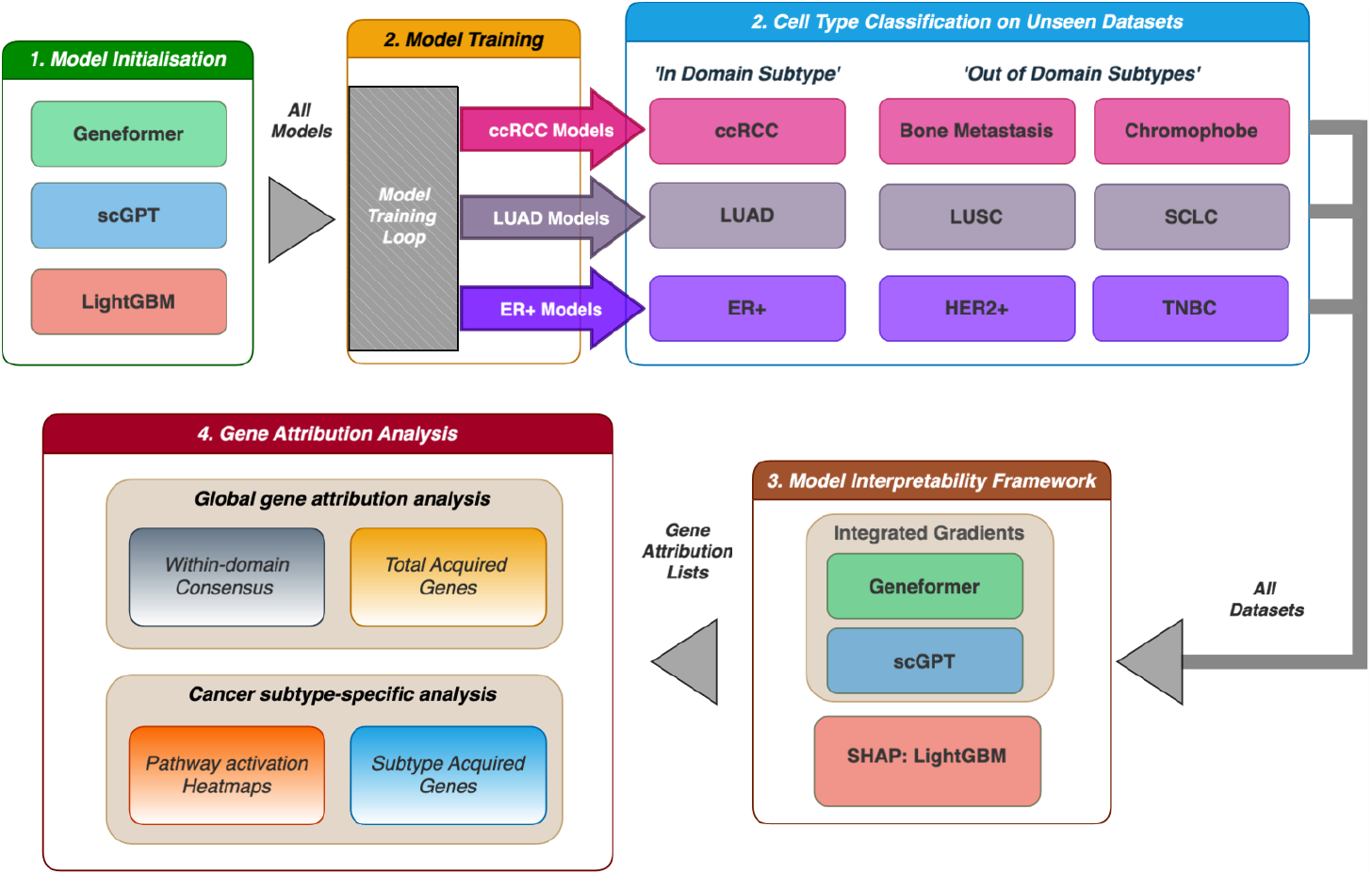
Pipeline schematic. Overview of the analytical pipeline. (A) Three models (Geneformer, scGPT, LightGBM) are initialised and fine-tuned on three cancer training domains: ccRCC, LUAD, and ER+ breast cancer. Trained models are evaluated on within-domain subtypes and out-of-domain rarer subtypes (Bone Metastasis, Chromophobe RCC; LUSC, SCLC; HER2+, TNBC). Integrated Gradients (for scFMs) and SHAP (for LightGBM) are applied to obtain gene attribution lists, which are analysed for within-domain consensus, acquired genes, pathway activation heatmaps, and subtype-specific gene acquisition.

## Results

### Study design

The overall study design is summarised in Figure 1. For each organ system, all models were trained on the common subtype and then evaluated on both matched validation cohorts and rarer subtypes. The task was binary cell-state classification: malignant tumour cells versus the most challenging normal comparator available in the target dataset, typically the tissue of origin or a closely related epithelial population. This design forced the models to distinguish malignant programmes from normal states that are transcriptionally similar to tumour cells rather than from easy immune-background negatives.

### Fine-tuned scFMs retain discrimination under clinically relevant subtype shift

Within domain, all three models performed strongly across the renal, lung, and breast settings (Figure 2A-C). In renal cancer, all three models achieved near-perfect discrimination on the external ccRCC cohorts: scGPT 0.983, 0.998 and 0.996 across the US, Lithuania and China datasets; Geneformer 0.980, 0.993 and 0.991; LightGBM 0.981, 0.997 and 0.994. This is consistent with the idea that simple supervised baselines remain hard to beat when train and test distributions are closely matched. Similar near-perfect discrimination was observed for the within-domain ER+ breast cohorts (scGPT 0.984 and 0.986; Geneformer 0.979 and 0.945; LightGBM 0.987 and 0.987) and for the LUAD metastatic cohorts closest to the training distribution (AUROC 0.96-1.00 for all models).

**Figure 2.**
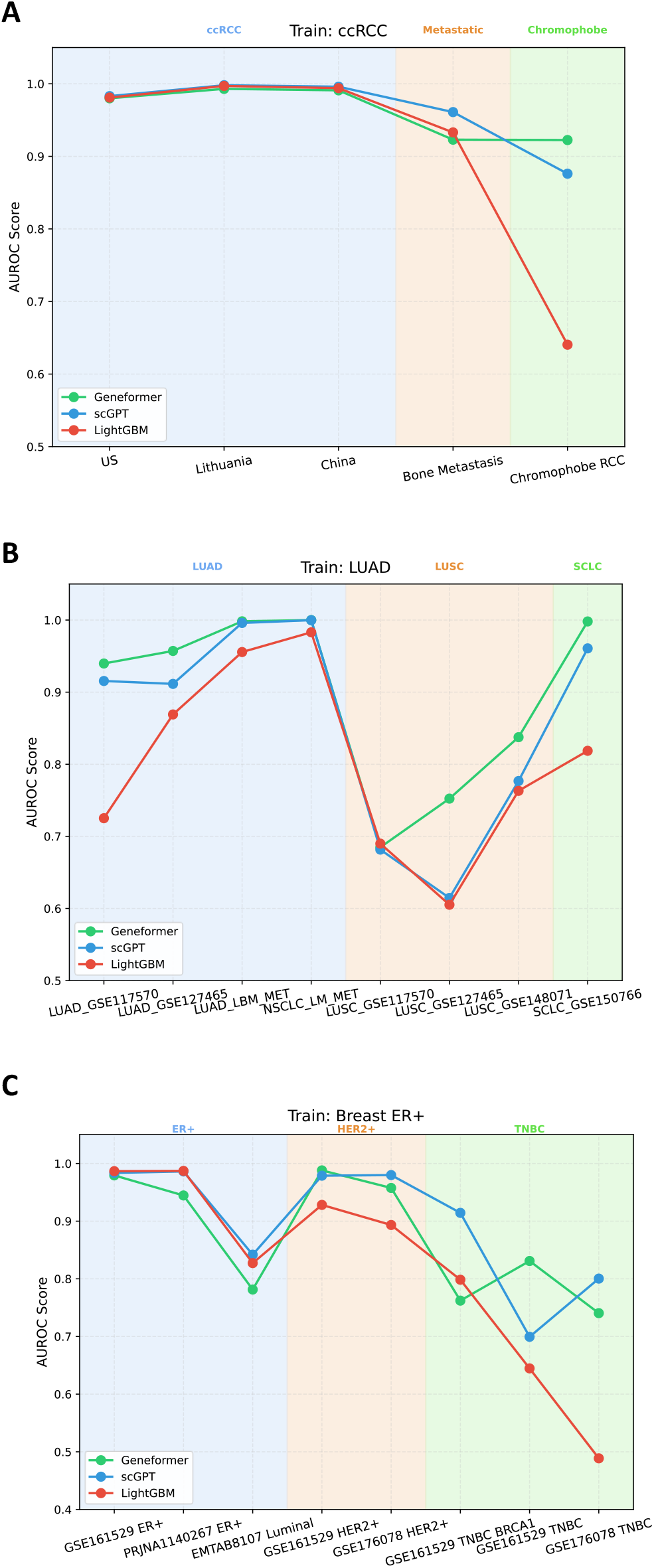
Cell type classification performance. (A) AUROC scores for ccRCC-trained models evaluated on within-domain ccRCC datasets (US, Lithuania, China) and out-of-domain Bone Metastasis and Chromophobe RCC datasets. (B) AUROC scores for LUAD-trained models evaluated on within-domain LUAD datasets and out-of-domain LUSC and SCLC datasets. (C) AUROC scores for ER+-trained models evaluated on within-domain ER+ datasets and out-of-domain HER2+, Luminal, and TNBC datasets. Foundation models generally outperform LightGBM on the most biologically distant out-of-domain tasks, although exceptions exist.

The more informative pattern emerged under out-of-domain transfer. In the renal setting, LightGBM remained competitive on bone metastasis (AUROC 0.933 versus 0.961 for scGPT and 0.923 for Geneformer) but collapsed on chromophobe RCC (AUROC 0.641), whereas scGPT achieved 0.876 and Geneformer 0.923 (Figure 2A). In the lung setting, all models struggled with LUSC, where AUROC ranged from 0.61 to 0.84 across cohorts with no model consistently dominant. However, on SCLC the foundation models were clearly superior: Geneformer reached 0.998, scGPT 0.961, and LightGBM 0.819 (Figure 2B). Notably, in the within-domain LUAD cohorts, LightGBM underperformed the foundation models on two of the four datasets (GSE117570: LightGBM 0.725 versus scGPT 0.916 and Geneformer 0.940; GSE127465: LightGBM 0.869 versus scGPT 0.911 and Geneformer 0.957), suggesting that even within-domain generalisation across distinct cohorts can be challenging for tree-based models.

In breast cancer, all models deteriorated on the broader luminal cohort (scGPT 0.842, Geneformer 0.781, LightGBM 0.827), but the foundation models generally showed smaller losses than LightGBM on HER2+ and TNBC cohorts (Figure 2C). On the GSE176078 HER2+ dataset, scGPT achieved 0.980, Geneformer 0.958 and LightGBM 0.893. The most striking divergence was on TNBC: on the GSE176078 TNBC cohort, scGPT achieved 0.800 and Geneformer 0.740, whereas LightGBM dropped to 0.489, essentially performing at chance. On the GSE161529 TNBC cohort, Geneformer (0.831) outperformed both scGPT (0.699) and LightGBM (0.645), while on the TNBC BRCA1 subset, scGPT led with 0.914 versus LightGBM at 0.798 and Geneformer at 0.762.

These results support a simple interpretation: LightGBM is excellent at interpolation within the learned decision boundary, whereas the foundation models are less fragile to changing context and are better able to preserve useful signal when both the malignant state and the normal comparator shift away from the original fine-tuning distribution. The advantage is not uniform across all scFMs or all subtypes, but the pattern is consistent: in the most biologically distant transfers (chromophobe RCC, SCLC, TNBC), the gap between foundation models and LightGBM is largest.

### Foundation models are context-sensitive, whereas LightGBM is attribution-rigid

To quantify how strongly each model changed its feature usage across datasets, we computed the mean pairwise Spearman correlation of the ranked attribution lists across all datasets (Figure 3A). LightGBM showed strikingly high stability in every organ system (renal 0.957, lung 0.965, breast 0.975), indicating that it largely reused the same ranked gene rules regardless of context. By contrast, scGPT showed intermediate stability (0.786, 0.826 and 0.862 across renal, lung and breast), whereas Geneformer was the most variable overall (0.777, 0.713 and 0.803).

**Figure 3.**
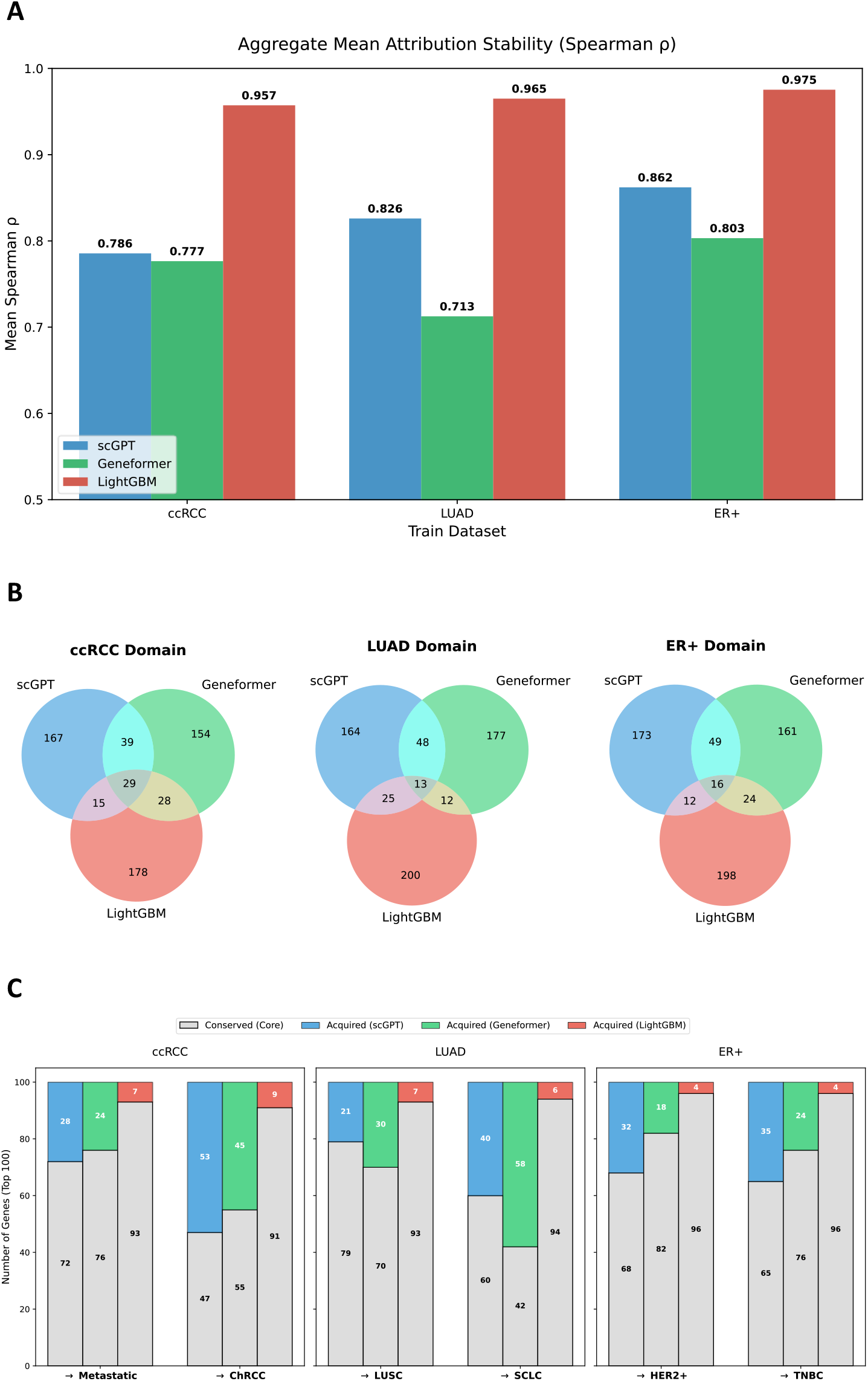
Attribution stability, within-domain consensus and gene acquisition under subtype shift. (A) Aggregate mean attribution stability (Spearman rho) for each model across the three cancer domains. LightGBM exhibits high stability (rho > 0.95) reflecting rigid gene usage, while both scGPT and Geneformer show lower stability (rho = 0.71-0.86) reflecting context-sensitive gene attribution. (B) Venn diagrams showing the overlap of the top 250 tumour-associated genes used within-domain by each model for ccRCC, LUAD, and ER+ domains. The triple intersection (29, 13, and 16 genes respectively) represents genes independently identified by all three models as important for each cancer type. (C) Stacked bar charts showing the composition of the top 100 attributed genes when shifting from within-domain to out-of-domain datasets. The conserved core (grey) represents genes retained from the within-domain consensus, while coloured segments represent genes newly acquired by each model. Foundation models acquire substantially more new genes than LightGBM. Together, these panels show that LightGBM maintains a near-fixed feature hierarchy across contexts, whereas the foundation models share a common gene core but flexibly recruit subtype-specific genes under domain shift.

**Figure 4.**
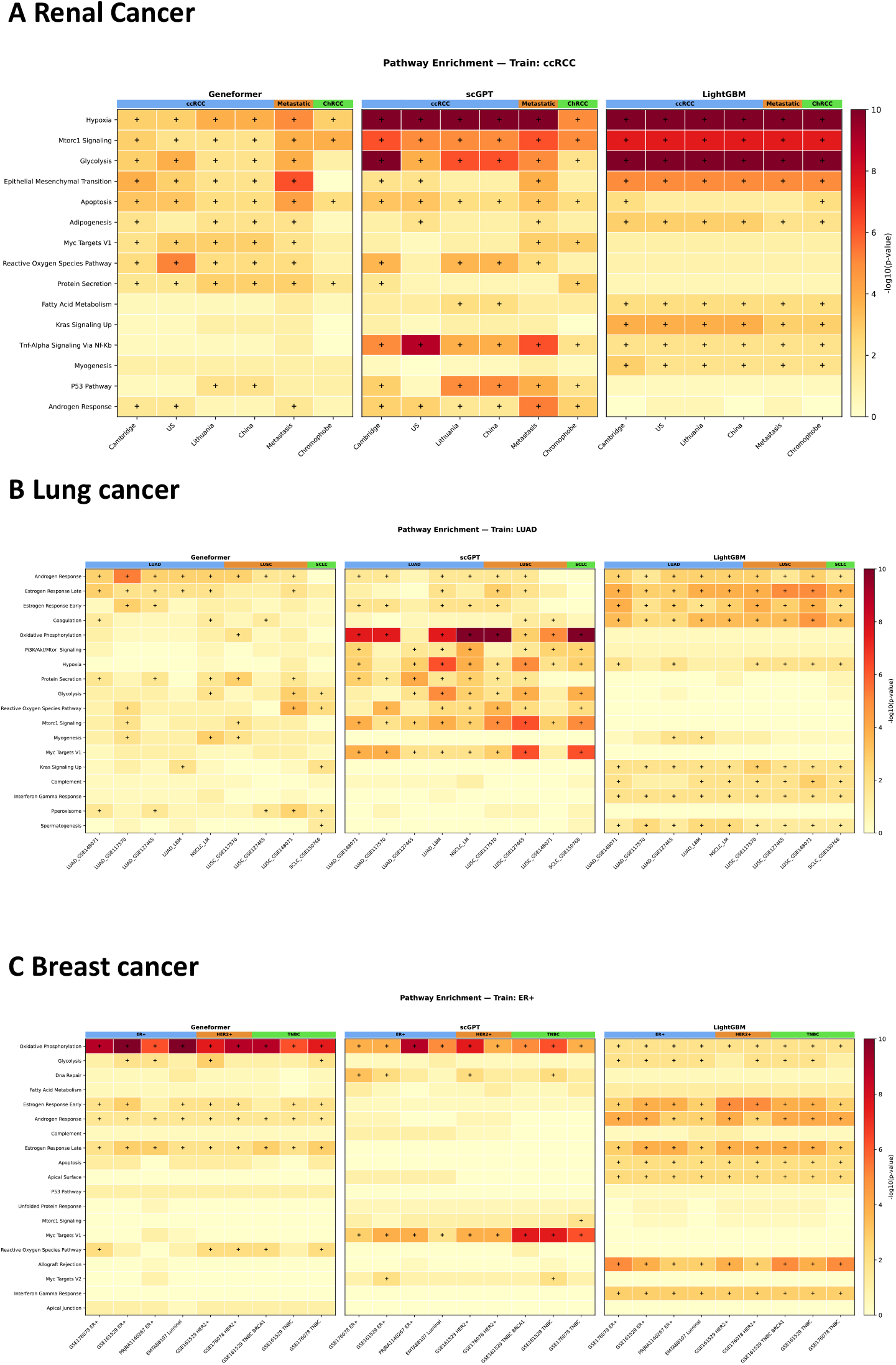
Pathway activation heatmaps. Pathway enrichment heatmaps for (A) Renal Cancer, (B) Lung Cancer, and (C) Breast Cancer. Each panel shows Geneformer, scGPT, and LightGBM side by side. Colour intensity represents −log10(p-value). Bold crosses indicate significant enrichment (P < 0.05) for a specific validation dataset. Foundation models show broader and stronger pathway activation in out-of-domain datasets compared to LightGBM.

This difference is consistent with model architecture. LightGBM applies fixed tree splits learned during training, so feature importance remains structurally constrained when the input distribution changes. Transformer-based scFMs, in contrast, condition each token representation on the broader transcriptomic context via the self-attention mechanism [3], allowing the apparent importance of a gene to shift depending on the cell state being analysed. The key question, therefore, is whether this context sensitivity reflects meaningful biological adaptation or merely unstable feature reuse?

### A shared gene consensus exists across model-specific solutions

Despite their different architectures, the three models converged on a gene consensus within each training domain (Figure 3B). The three-way intersection contained 29 genes in ccRCC, 13 in LUAD and 16 in ER+ breast cancer, suggesting that correct within-domain classification is supported by a limited set of canonical disease-associated features.

At the same time, the pairwise-only overlaps and model-specific regions were large. This indicates that similar classification performance does not imply identical internal solutions. The models agree on a compact common programme but diverge in how much additional signal they recruit around that core. In practice, this makes interpretability essential: high AUROC alone cannot reveal whether two models arrive at a correct answer through the same biology or through different, only partially overlapping feature sets.

### Rare-subtype transfer is accompanied by acquisition of new genes

We next asked how much each model’s top-ranked tumour-supporting attributed genes were conserved versus newly acquired under subtype shift (Figure 3C). Across every out-of-domain comparison, LightGBM retained the overwhelming majority of its top 100 genes, keeping 91-96 genes from the original within-domain set and adding only 4-9 new genes. The foundation models behaved very differently. In ccRCC to chromophobe RCC transfer, scGPT and Geneformer acquired 53 and 45 new genes, respectively, versus only 9 for LightGBM. In LUAD to SCLC, the shift was even stronger, with 40 acquired genes for scGPT and 58 for Geneformer, compared with 6 for LightGBM. In ER+ to HER2+ or TNBC, the foundation models again incorporated broader rare-subtype-specific programmes than the baseline.

This pattern argues against the idea that foundation models succeed only by reusing a frozen pre-training representation. Instead, fine-tuning appears to establish a transferable cancer prior that can be selectively extended when the model encounters a related but distinct disease context.

### Acquired genes are biologically interpretable and differ by model

The rank-shift plots for individual domain transitions illustrate how these adaptive programmes manifest at the gene level (Figures 5-7). In renal cancer, scGPT prominently upweighted genes such as SOD2, MIF and BIRC3 in bone metastasis, while Geneformer acquired a distinct set including C3, MT-ND5 and SLPI (Figure 5A). To assess whether these acquired genes carry functional relevance beyond attribution ranking, we examined their association with patient survival using TCGA data. Two acquired bone metastasis genes, SAA1 (scGPT-acquired) and SLPI (Geneformer-acquired), were significantly associated with poor overall survival in renal cell carcinoma (SAA1: log-rank P = 1.18 × 10^-9, HR = 2.47; SLPI: log-rank P = 1.02 × 10^-6, HR = 2.04), whereas two scGPT-dropped genes, SF3B6 and RACK1, showed no significant survival association (Figure 5B-C). In lung cancer, scGPT acquired canonical squamous-associated markers including KRT5, KRT17 and GPX2 when transferring from LUAD to LUSC, while dropping adenocarcinoma markers such as AGR2, MUC1 and KRT7 (Figure 6A). Differential expression analysis confirmed that KRT5 is significantly overexpressed in LUSC but underexpressed in LUAD, while AGR2 shows the opposite pattern, validating that the model’s gene shifts mirror genuine subtype-specific biology (Figure 6B-C). In breast cancer, the ER+ to TNBC shift was accompanied by acquisition of spliceosome-associated genes including SNRPG, TRA2B and SNRPD1, and loss of ER-associated genes such as GATA3 and FOXA1 (Figure 7A). Subtype-stratified expression analysis confirmed that SNRPG expression increases progressively from normal tissue to TNBC (Luminal vs TNBC P = 3.23 × 10^-13), whereas FOXA1 is highly expressed in luminal tumours but near-absent in TNBC (Luminal vs TNBC P < 10^-12), consistent with the direction of scGPT’s gene reweighting (Figure 7B-C).

**Figure 5.**
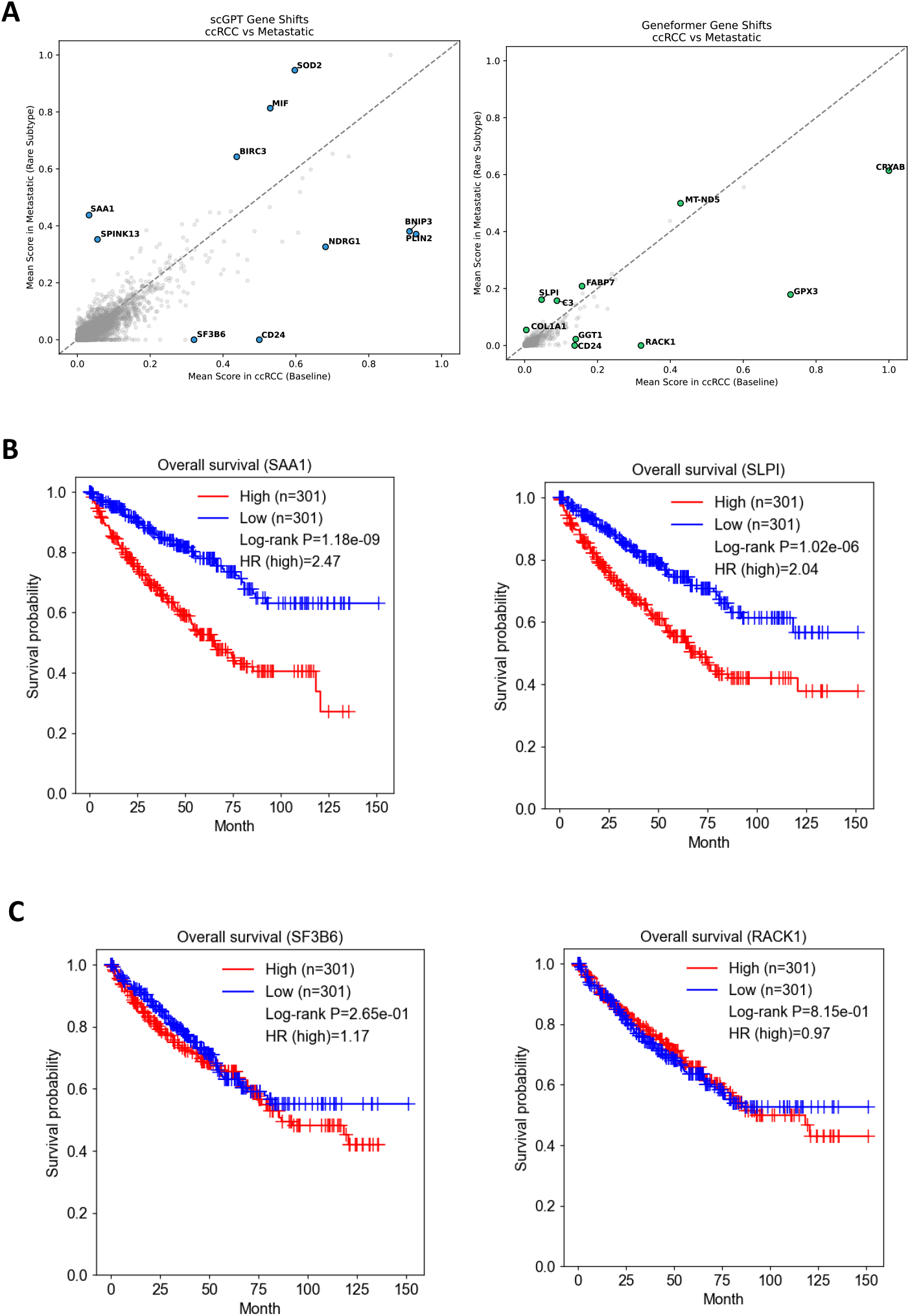
Gene-rank shifts in renal cancer transfer and survival validation of acquired genes. (A) Rank-shift scatter plots for scGPT (left) and Geneformer (right) showing gene importance changes when transferring from ccRCC to bone metastasis. Genes above the diagonal gained importance in the metastatic context (acquired), while genes below the diagonal lost importance (dropped). scGPT acquires biologically relevant genes including SOD2, MIF, BIRC3, SAA1 and NDRG1, while dropping SF3B6 and CD24. Geneformer acquires CRYAB, MT-ND5 and SLPI, while dropping RACK1, GGT1 and CD24. (B) Overall survival analysis for two acquired genes, SAA1 (scGPT-acquired) (log-rank P = 1.18 × 10^-9, HR = 2.47) and SLPI (Geneformer-acquired; log-rank P = 1.02 × 10^-6, HR = 2.04), showing that high expression of both genes is significantly associated with poor prognosis in renal cell carcinoma. (C) Overall survival analysis for two scGPT-dropped genes, SF3B6 (log-rank P = 0.265, HR = 1.17) and RACK1 (log-rank P = 0.815, HR = 0.97), showing no significant survival association. These results indicate that genes acquired by the foundation models under domain shift carry prognostic relevance, whereas genes that lose importance do not. Survival analyses were performed using scCancerExplorer [29].

**Figure 6.**
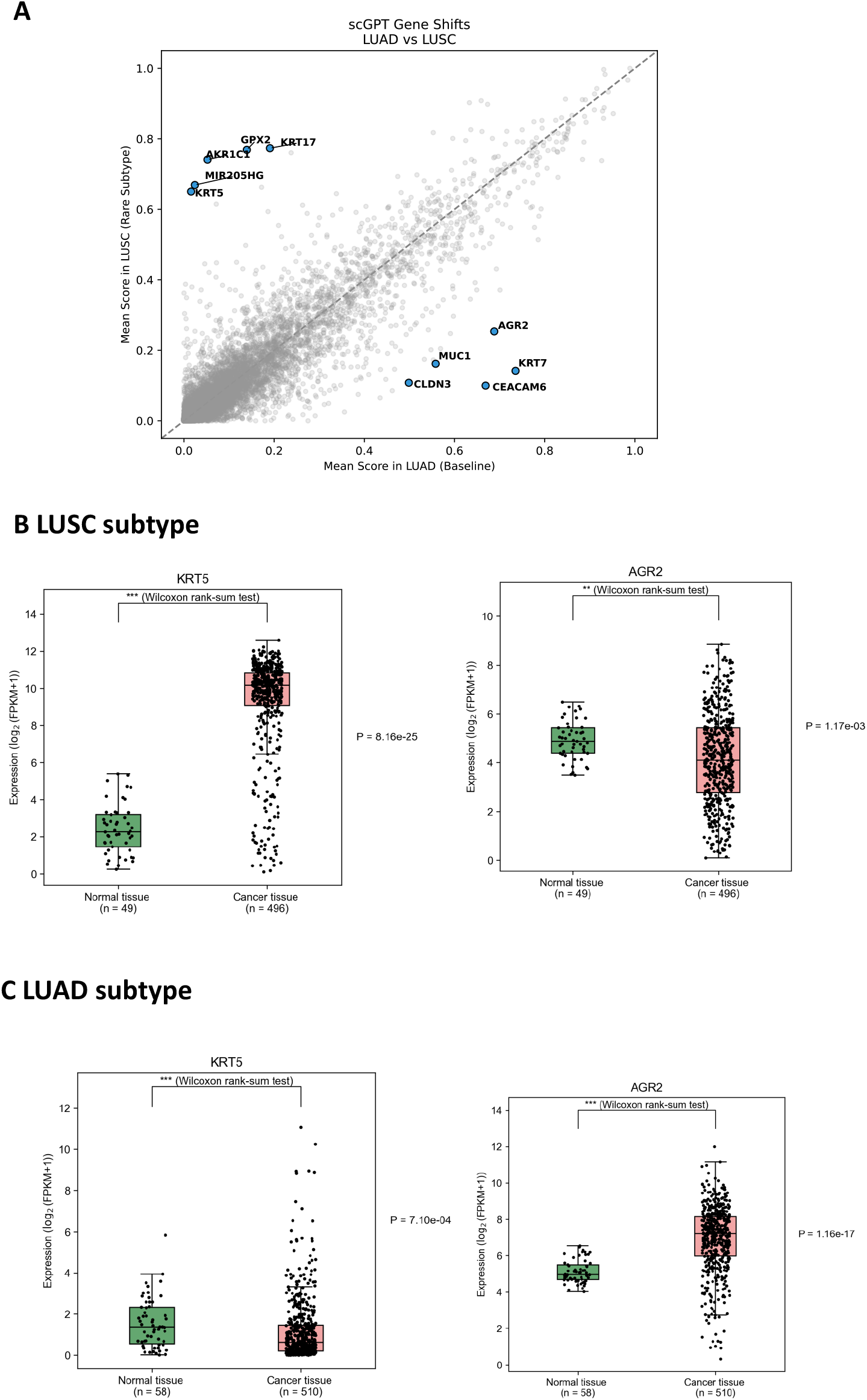
Gene-rank shifts in lung cancer transfer and differential expression validation. (A) Rank-shift scatter plot for scGPT showing gene importance changes when transferring from LUAD to LUSC. Genes above the diagonal gained importance in the squamous context (acquired), while genes below the diagonal lost importance (dropped). scGPT acquires squamous-associated markers including KRT17, KRT5, GPX2, AKR1C1 and MIR205HG, while dropping adenocarcinoma markers AGR2, MUC1, CLDN3, KRT7 and CEACAM6. (B) UALCAN differential expression analysis in LUSC, showing that KRT5 is significantly overexpressed in cancer tissue relative to normal tissue (P = 8.16 × 10^-25), whereas AGR2 shows only modest differential expression (P = 1.17 × 10^-3). (C) UALCAN differential expression analysis in LUAD, showing that KRT5 is significantly underexpressed in cancer tissue (P = 7.10 × 10^-4), while AGR2 is significantly overexpressed (P = 1.16 × 10^-17). The reversal of KRT5 and AGR2 expression patterns between LUSC and LUAD confirms that scGPT’s gene acquisition and loss under domain shift reflects genuine subtype-specific biology rather than arbitrary feature drift. Differential expression analyses were performed using the UALCAN portal [27, 28].

**Figure 7.**
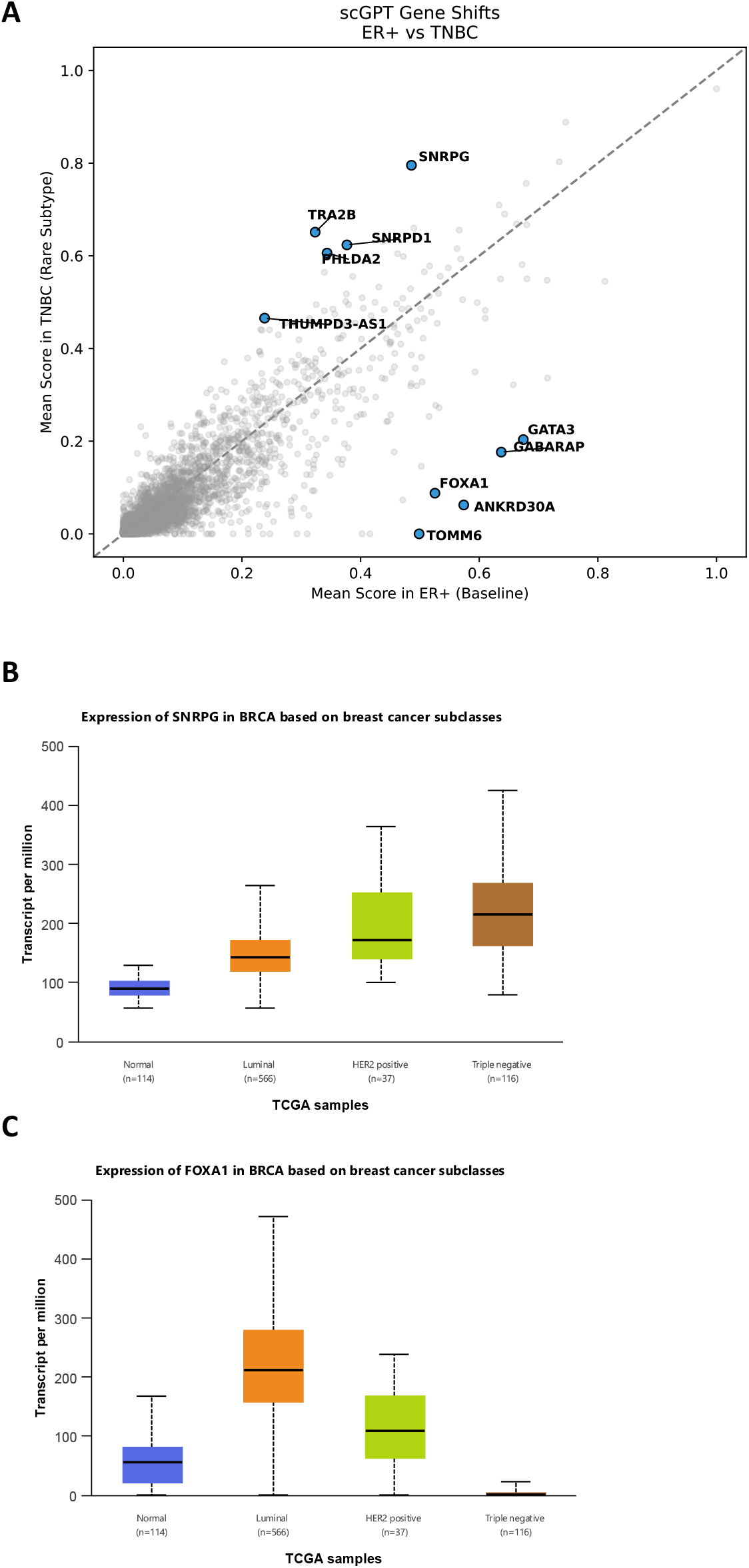
Gene-rank shifts in breast cancer transfer and subtype-specific expression validation. (A) Rank-shift scatter plot for scGPT showing gene importance changes when transferring from ER+ to TNBC. Genes above the diagonal gained importance in the TNBC context (acquired), including SNRPG, TRA2B, SNRPD1, PHLDA2 and THUMPD3-AS1, while genes below the diagonal lost importance (dropped), including GATA3, GABARAP, FOXA1, ANKRD30A and TOMM6. (B) UALCAN expression of SNRPG across TCGA breast cancer subclasses. SNRPG expression increases progressively from normal tissue through luminal and HER2-positive to triple-negative breast cancer (Luminal vs TNBC P = 3.23 × 10^-13; Normal vs TNBC P < 10^-12), consistent with scGPT acquiring this gene when shifting to the TNBC context. (C) UALCAN expression of FOXA1 across TCGA breast cancer subclasses. FOXA1 is highly expressed in luminal tumours but near-absent in TNBC (Luminal vs TNBC P < 10^-12), consistent with scGPT dropping this gene under ER+ to TNBC transfer. These expression patterns confirm that scGPT’s gene acquisition and loss reflect the known molecular divergence between ER+/luminal and triple-negative breast cancer. Expression analyses were performed using the UALCAN portal [27,28].

These examples emphasise that foundation-model flexibility is structured rather than arbitrary. The models do not simply reshuffle all genes under shift; they selectively elevate genes consistent with the biology of the target subtype, and external validation through survival and differential expression analyses confirms that acquired genes carry measurable clinical and biological relevance. Notably, the rank-shift plots in the lung and breast settings focus on scGPT because it exhibited the most visually distinct gene-level shifts in those transitions. Geneformer acquired substantial numbers of new genes, as quantified in Figure 3C, but its rank-shift patterns were less visually separated among the highest-ranked genes in those settings.

### Pathway analysis supports biologically grounded adaptation

Gene-level variation becomes even more interpretable at the pathway level. Hallmark enrichment on the top attributed genes revealed that the models recurrently activated biologically plausible cancer programmes across organs, including hypoxia, glycolysis, epithelial-mesenchymal transition, oxidative phosphorylation, MYC signalling and inflammatory responses (Figure 4). Importantly, the heatmaps show both conservation and adaptation: some pathways are maintained across all datasets within an organ, whereas others intensify or appear only in the rare subtypes. Notably, the Hypoxia pathway is consistent with the well-characterised VHL/HIF axis in ccRCC [21].

The functional relevance of these context-dependent pathway activations is particularly evident in the rare-subtype transfers. In SCLC, both Geneformer and scGPT strongly activated the Reactive Oxygen Species and Glycolysis pathways, neither of which was enriched in the LightGBM attribution lists for the same dataset (Figure 4B). scGPT additionally repurposed Oxidative Phosphorylation and Myc Targets across the lung datasets, pathways that LightGBM did not engage at all. These observations suggest that the foundation models recruit metabolic and proliferative programmes that are biologically appropriate for the target subtype but invisible to a classifier constrained by a fixed feature hierarchy.

The breast cancer analysis provides a particularly clear example. Oxidative phosphorylation was repeatedly recovered across all models, but the scFMs encoded that pathway with different genes and with different degrees of dataset dependence (Figure 8). LightGBM relies on a smaller fixed panel, Geneformer distributed importance over a broader and more redundant set, and scGPT showed a sparser, more context-specific pattern. Thus, pathway-level agreement can coexist with gene-level divergence, suggesting that the models may represent the same biological process at different levels of abstraction.

**Figure 8.**
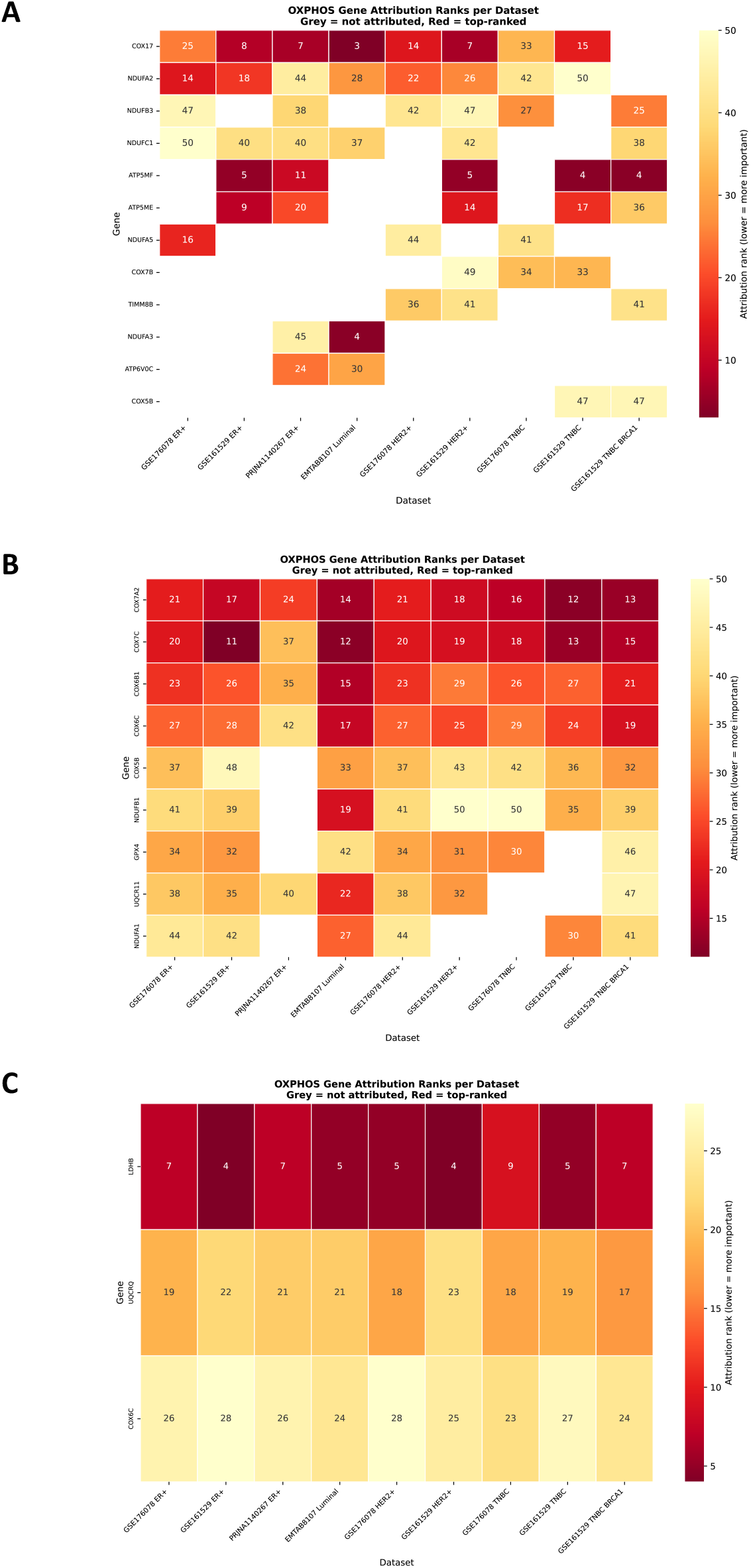
Context-specific encoding of oxidative phosphorylation in breast cancer. Heatmaps showing OXPHOS gene attribution ranks per breast cancer dataset for (A) scGPT, (B) Geneformer, and (C) LightGBM. Foundation models use different OXPHOS genes depending on the dataset context, while LightGBM shows a rigid, uniform pattern

### Consensus foundation model genes are multi-omically validated in ccRCC

To move beyond attribution ranking and assess whether the genes jointly identified by the foundation models carry independent biological and clinical support, we took the intersection of the scGPT and Geneformer within-domain attributed gene lists for ccRCC, excluding genes that were also top-ranked by LightGBM, and grouped the resulting set by biological mechanism. We selected a representative gene from each of the four most prominent mechanistic categories and confirmed that all four were consistently upregulated in tumour cells across the four single-cell RNA-seq ccRCC datasets (training, US, Lithuania and China cohorts).

We then examined each representative gene using bulk TCGA RNA expression, DNA promoter methylation and overall survival data (Figure 9). EIF4EBP1, representing the hypoxia / HIF-adaptation / glycolysis / redox defence programme, was significantly overexpressed in tumour tissue (P = 4.25 × 10^-30), showed altered promoter methylation (P = 5.29 × 10^-13) and was associated with poor survival (HR = 1.74, P = 1.29 × 10^-4). TMEM258, representing ER proteostasis / folding / translocation / glycosylation, was similarly overexpressed (P = 2.13 × 10^-19) with significant methylation change (P = 3.42 × 10^-4) and a modest survival association (HR = 1.33, P = 4.64 × 10^-2). CD63, representing lysosome / protease control / nutrient sensing / invasion trafficking, showed strong overexpression (P = 3.56 × 10^-37), altered methylation (P = 2.12 × 10^-3) and significant survival association (HR = 1.40, P = 1.92 × 10^-2). YBX3, representing signalling / transcriptional control / RNA regulation, was overexpressed (P = 2.58 × 10^-36) with highly significant methylation change (P = 1.00 × 10^-9) and poor survival association (HR = 1.44, P = 1.05 × 10^-2).

**Figure 9.**
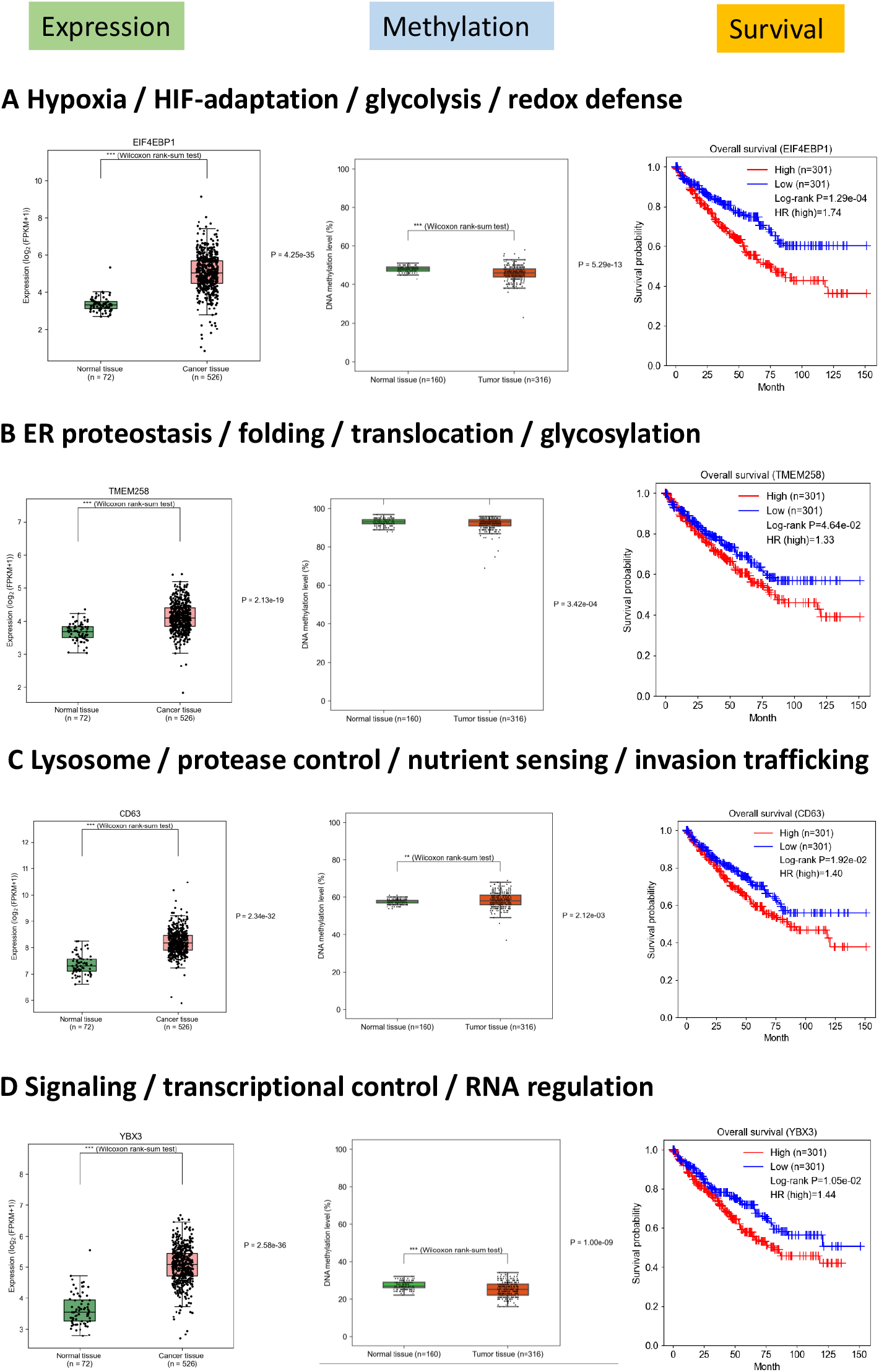
Multi-omic validation of consensus foundation model genes in ccRCC. Genes from the intersection of scGPT and Geneformer within-domain attributed gene lists (excluding LightGBM) were grouped by biological mechanism. All representative genes shown were consistently upregulated in tumour cells across four single-cell RNA-seq ccRCC datasets (training, US, Lithuania and China cohorts). For each gene, RNA expression (left), DNA promoter methylation (centre) and overall survival (right) are shown using TCGA data from scCancerExplorer [29]. (A) Hypoxia / HIF-adaptation / glycolysis / redox defence, represented by EIF4EBP1 (expression P = 4.25 × 10^-30; methylation P = 5.29 × 10^-13; survival log-rank P = 1.29 × 10^-4, HR = 1.74). (B) ER proteostasis / folding / translocation / glycosylation, represented by TMEM258 (expression P = 2.13 × 10^-19; methylation P = 3.42 × 10^-4; survival log-rank P = 4.64 × 10^-2, HR = 1.33). (C) Lysosome / protease control / nutrient sensing / invasion trafficking, represented by CD63 (expression P = 3.56 × 10^-37; methylation P = 2.12 × 10^-3; survival log-rank P = 1.92 × 10^-2, HR = 1.40). (D) Signalling / transcriptional control / RNA regulation, represented by YBX3 (expression P = 2.58 × 10^-36; methylation P = 1.00 × 10^-9; survival log-rank P = 1.05 × 10^-2, HR = 1.44). All four representative genes show significant overexpression in tumour tissue, altered promoter methylation and association with poor overall survival, supporting the biological and clinical relevance of the foundation model consensus gene set.

These results demonstrate that genes prioritised by the consensus of two independent foundation models, but not by LightGBM, are not merely correlated transcriptomic markers. They show convergent evidence across expression, epigenetic regulation and patient outcome, supporting the idea that the foundation models capture biologically grounded tumour programmes that a rigid tree-based classifier does not prioritise.

## Discussion

This study addresses a practical question in cancer single-cell analysis: when does the extra complexity of a foundation model become useful? Our results suggest a clear answer. When applied to datasets within domain, strong classical baselines such as LightGBM remain difficult to outperform. Under biologically meaningful subtype shift, however, the fine-tuned scFMs were generally more robust. Their advantage was not universal and depended on the organ system and target subtype, but the pattern was consistent enough to support the idea that pre-training provides transferable structure that becomes most valuable when the downstream task moves away from the original supervised distribution, addressing a critical bottleneck in the application of AI to rare diseases [14].

The interpretability analyses were essential for understanding why. LightGBM’s strength is also its limitation: it retains a very stable feature hierarchy, which is advantageous when the target data obey the same rules but leaves little room for adaptation when disease biology changes. The foundation models behaved differently. Their gene-level attributions were less stable, but this instability was accompanied by coherent pathway reuse and by subtype-appropriate acquisition of new genes. In other words, the same flexibility that makes scFMs harder to summarise with a single static feature list appears to underlie their stronger transfer behaviour. This aligns with recent benchmarking efforts demonstrating that fine-tuning significantly enhances the adaptability of scFMs compared to zero-shot applications [9, 10].

The biological relevance of this adaptation is supported by multiple lines of evidence. First, the within-domain consensus genes across all three models correspond to classical biomarkers for each cancer type. Second, the acquired genes in out-of-domain settings correspond to known biology: for example, scGPT drops ER-associated genes (GATA3, FOXA1) and acquires new markers when shifting from ER+ to TNBC, consistent with the distinct molecular profiles of these subtypes [22]. Third, pathway enrichment analysis confirms that model-attributed genes are enriched for cancer hallmark pathways relevant to the specific cancer subtype, such as the VHL/HIF-driven hypoxia response in ccRCC [22, 23].

The two scFMs were not interchangeable. scGPT often produced the clearest gene-level shifts and the most consistent breast cancer transfer (AUROC 0.800 on GSE176078 TNBC versus Geneformer 0.740), whereas Geneformer was the stronger model on SCLC (0.998 versus 0.961) and on the GSE161529 TNBC cohort (0.831 versus 0.699). Geneformer also frequently distributed importance across a broader set of pathway members. This suggests that there may not be a single best scFM for all cancer-transfer tasks. Instead, different pre-training objectives and tokenisation strategies may favour different modes of adaptation.

Taken together, the results support a practical framework for deploying scFMs in oncology. Foundation models should not be treated as universal drop-in replacements for strong baselines, especially not in zero-shot settings [6]. Their value emerges most clearly when they are lightly adapted to a disease context and then challenged with related but data-poor clinical subtypes. In those settings, interpretability is not a cosmetic add-on; it is the mechanism that allows us to distinguish adaptive reuse of transferable biology from opaque overfitting.

### Limitations and future directions

Several limitations should be acknowledged. First, the IG analyses used a padding-token-based baseline, and alternative baselines could alter the exact attribution values [18]. The choice of baseline is a known challenge in interpretable machine learning for biology [19], and it remains an open question whether the pad token represents a truly neutral reference point. Second, our primary comparisons focused on positively tumour-supporting attributed genes; negatively attributed features that support the normal prediction were not analysed in depth. Third, LightGBM was the only classical baseline included, so the results should not be overgeneralised to all non-foundation approaches. Finally, attribution and enrichment provide mechanistic clues but do not establish causality. Experimental validation of newly acquired rare-subtype genes remains an important next step.

## Methods

### Task definition and evaluation framework

Three organ-specific training settings were considered: renal (train: ccRCC), lung (train: LUAD) and breast (train: ER+). For each setting, models were evaluated on both matched validation cohorts of the same subtype and on rarer out-of-domain subtypes. The prediction task was binary cell-state classification, with tumour cells as the positive class and a challenging normal comparator as the negative class. In each validation cohort, the normal class was chosen to reflect the tissue of origin whenever possible, thereby making the task harder than discrimination against immune-background cells alone. Class sizes were balanced during training and evaluation, and AUROC was used as the primary performance metric.

### Models and fine-tuning

We evaluated Geneformer [2], scGPT [1] and LightGBM [11]. Both Geneformer and scGPT were accessed via the helical package [24] where we used the model versions trained predominantly on non-diseased tissues [1, 2]. All three models were fine-tuned or trained on the common subtype for each organ and then applied without retraining to the corresponding validation cohorts. LightGBM served as a strong supervised baseline because it is efficient, interpretable and often highly competitive on cell-state classification. The transformer models were used in a task-adapted setting rather than a zero-shot setting because the goal of the study was to measure domain adaptation after limited cancer-specific supervision rather than to benchmark frozen embeddings [6].

### Preprocessing

Preprocessing was intentionally kept minimal for the foundation models so that evaluation reflected transfer from the pre-trained representation with limited task-specific adaptation. LightGBM and scGPT were trained on 10,000 highly variable genes (HVGs) from each training dataset, whereas Geneformer was trained on all genes due to its rank-based encoding. For LightGBM, expression matrices were scaled and log-normalised before model fitting.

### Integrated gradients and SHAP attributions

To identify the genes that most strongly supported malignant predictions, we applied Integrated Gradients (IG) to the fine-tuned scFMs [18]. IG returns signed attributions for each input feature relative to a chosen baseline, allowing genes that push predictions toward tumour or normal states to be separated. Attributions were computed per cell and then summed over 500 randomly selected tumour cells from each dataset. In exploratory analyses, increasing the number of cells beyond 500 had little impact on the aggregated rankings. We also experimented with mean attribution values, dividing each gene’s attribution by the number of cells in which it appeared, but this yielded fewer significantly enriched pathways owing to a bias toward genes that appear less regularly rather than globally important ones.

Because LightGBM is tree-based, directional feature attributions were obtained using SHAP values [20]. For cross-model comparisons focused on tumour-supporting signal, negatively attributed IG genes and zero-importance SHAP genes were excluded, and the remaining importance values were min-max normalised within each gene list.

### Attribution stability and consensus analyses

Across all datasets within each organ system, ranked gene lists were compared pairwise using Spearman rank correlation. The mean of these pairwise correlations provided a measure of attribution stability for each model. To identify a within-domain consensus, we combined the training dataset with matched validation datasets from the same subtype and computed a consensus rank for each model. The top 250 tumour-associated genes per model were retained, and intersections between models were visualised to distinguish shared versus model-specific signal.

### Domain-shift acquisition analysis

For each out-of-domain subtype, top-ranked genes were divided into a conserved core (genes also present in the corresponding within-domain list) and an acquired set (genes present only after subtype shift). Rank-shift scatter plots were then used to visualise which genes gained, retained or lost importance relative to the baseline training subtype.

### Pathway enrichment analysis

Biological interpretation of the attributed gene lists was performed on the top 50 tumour-associated genes per dataset using GSEApy [25], querying the MSigDB Hallmark collection [26]. For each organ system, enriched pathways from the three models were combined into a shared axis so that pathway usage could be compared side by side. Cells marked with a plus sign in the heatmaps indicate pathway enrichment at nominal P < 0.05 in the corresponding validation dataset.

## Data Availability

All datasets analysed during this study are publicly available. Where applicable, Gene Expression Omnibus (GEO) identifiers are provided. Multiple datasets were reused by sub-setting into the different patient subtypes.

### Renal Cancer

The clear cell renal cell carcinoma (ccRCC) training dataset is available on Mendeley Data (https://data.mendeley.com/datasets/g67bkbnhhg/1). Additional ccRCC validation datasets were obtained from GEO under accession numbers GSE159115 (US cohort), GSE242299 (Lithuania cohort), and GSE156632 (China cohort). The Chromophobe RCC dataset was subset from the US cohort (GSE159115). The bone metastasis dataset is available under accession number GSE202813.

### Lung Cancer

All lung tumour datasets were accessed via the Tumor Immune Single-cell Hub (TISCH) database (https://tisch.compbio.cn/gallery/?cancer=NSCLC&cancer=SCLC&celltype=&species=). The normal class validation datasets (Basal for LUSC, Alveolar Epithelial Type 2 for others) were obtained from the Human Lung Cancer Atlas and downloaded as an h5ad file (https://explore.data.humancellatlas.org/projects/6936da41-3692-46bb-bca1-cd0f507991e9). For Lung Adenocarcinoma (LUAD), the training dataset is available under GSE148071. Validation datasets include GSE127465, GSE117570, GSE143423 (late brain metastasis), and GSE150660 (leptomeningeal metastases). For Lung Squamous Cell Carcinoma (LUSC), datasets GSE148071, GSE127465, and GSE117570 were utilised. The Small Cell Lung Cancer (SCLC) dataset is available under GSE150766.

### Breast Cancer

The majority of breast cancer datasets were downloaded from the TISCH database (https://tisch.compbio.cn/gallery/?cancer=BRCA&species=). When normal epithelial cells were not available, subtype-specific normal epithelial cells from GSE161529 were used. For the ER+ subtype, the training dataset is available under GSE176078. Additional datasets include GSE161529, PRJNA1140267 (downloaded from Zenodo: https://zenodo.org/records/13743374), and EMTAB8107 (Luminal subtype, downloaded from the original publication). For the HER2+ subtype, datasets GSE161529 and GSE176078 were used. For Triple-Negative Breast Cancer (TNBC), datasets GSE161529 (BRCA1 subset), GSE161529 (non-BRCA1 subset), and GSE176078 were analysed.

## Code Availability

All code used in this study is publicly available at https://github.com/jameswallacew8/DomainAdaptation.git. This includes scripts for model fine-tuning, cell type classification, integrated gradients attribution analysis, SHAP analysis, and pathway enrichment.

## Contributions

JW conducted the data analysis. G.Y. conceptualised the project. JW and GY drafted the first version of the manuscript. G.Y. and N.H. provided overall guidance. All authors reviewed the manuscript.

## Acknowledgements

We thank Mac Walker for his contributions to the data analysis.

## Notes

### Competing Interest Statement

The authors have declared no competing interest.

